# Tyrosyl-DNA phosphodiesterase 1 and topoisomerase I activities as predictive indicators for Glioblastoma susceptibility to genotoxic agents

**DOI:** 10.1101/700039

**Authors:** Wenjie Wang, Monica Rodriguez-Silva, Arlet M. Acanda de la Rocha, Aizik L. Wolf, Yanhao Lai, Yuan Liu, William C. Reinhold, Yves Pommier, Jeremy W. Chambers, Yuk-Ching Tse-Dinh

## Abstract

**Background:** Glioblastoma (GBM) patients have an estimated survival of ∼15 months with treatment, and the standard of care only modestly enhances patient survival. Identifying biomarkers representing vulnerabilities may allow for selection of efficacious chemotherapy options to address personalized variations in GBM tumors. Irinotecan, currently in clinical trials for GBM, targets topoisomerase I (TOP1) by forming a ternary DNA-TOP1 cleavage complex (TOP1cc) inducing apoptosis. Tyrosyl-DNA phosphodiesterase 1 (TDP1) is a crucial repair enzyme that may reduce the effectiveness of irinotecan.

**Methods:** We treated GBM cell lines with increasing concentrations of irinotecan and compared the IC_50_ values. TOP1 and TDP1 protein levels from each cell type as well as GBM patient tumors were determined by Western blot analysis, while activity levels were ascertained by specific enzymatic assays. Cellular TDP1 was elevated by ectopic expression of wild-type or mutant TDP1.

**Results:** After comparing cellular susceptibility to TDP1 and TOP1 concentrations and activities, we found that the TDP1/TOP1 activity ratio had the strongest correlation (Pearson correlation coefficient R = 0.92) with IC_50_ values following irinotecan treatment. Increasing the TDP1/TOP1 activity ratio by ectopic expression of wild-type TDP1 increased in irinotecan IC_50_, while expression of the TDP1 catalytic-null mutant did not alter the susceptibility to irinotecan.

**Conclusions:** TDP1/TOP1 activity ratio may be a new predictive indicator for GBM vulnerability to irinotecan, allowing for selection of individual patients for irinotecan treatment based on risk-benefit. Moreover, TDP1 inhibitors may be a novel combination treatment with irinotecan to improve GBM patient responsiveness to genotoxic chemotherapies.

**Key Points:**

1. TDP1/TOP1 activity ratio correlates with irinotecan sensitivity in GBM cell lines.
2. TDP1 and TOP1 protein levels are not reliable predictors for irinotecan activity.
3. TDP1 inhibition is a plausible approach to improve irinotecan effectiveness in GBM.

**Importance of the Study:** The current standard of care (surgery, radiation, and chemotherapy) for GBM patients modestly enhances survival beyond ∼15 months. Thus, there is a great need for effective therapies and biomarkers that address personalized variations in GBM tumors to improve treatment outcome. Topoisomerase I (TOP1) is the target of irinotecan. The repair enzyme tyrosyl-DNA phosphodiesterase 1 (TDP1) is known to excise irinotecan-induced TOP1-DNA cleavage complexes (TOP1ccs). Consequently, this study examines the relationship between TOP1 and TDP1 expression and activities in GBM cells and their correlation with irinotecan sensitivity. The results reveal that the TDP1/TOP1 activity ratio predicts irinotecan vulnerability in GBM cell lines. TDP1/TOP1 activity ratio was found to vary among GBM patient tumors. This potential predictive indicator may permit selection of patients responsive to irinotecan based on the capacity to repair TOP1cc. Moreover, inhibitors of TDP1 may represent a promising approach to enhance irinotecan efficacy in GBM patients.

## Introduction

Glioblastoma (GBM) is the most common and aggressive primary brain tumor in adults with a dismal prognosis indicated by a 14.6-month median survival after surgical resection, radiotherapy, and chemotherapy treatment with temozolomide (TMZ).^1^ Irinotecan is used to treat metastatic colorectal cancer and small cell lung cancer (SCLC).^2, 3^ Irinotecan has some capacity to cross the blood-brain barrier,^4^ and was shown to be effective against recurrent malignant glioma in early phase II clinical trials.^5, 6^ However, other clinical trials could not confirm the activity from irinotecan monotherapy.^7, 8^ Studies of irinotecan in combination therapy have yielded modest activity as well.^7, 9–11^ These seemingly discrepant results may indicate an underlying variability in the susceptibility of patients in each trial towards the genotoxic chemotherapy. Thus, to improve clinical outcomes and minimize toxic side-effects, identifying biomarkers for GBM susceptibility to irinotecan would be instrumental for selecting patients for irinotecan treatment.

Irinotecan, its active metabolite SN-38, and topotecan, are chemotherapeutic camptothecin derivatives (CPTs) acting as topoisomerase I (TOP1) poisons.^12^ Human TOP1 is vital for replication and transcription, as it relaxes DNA superhelical tension through phosphodiester bond cleavage, strand rotation, and DNA resealing steps.^13^ During the cleavage step, TOP1 covalently attaches to the 3’-end of DNA with its catalytic tyrosine residue forming reversible DNA-TOP1 cleavage complexes (TOP1cc). Stabilization of the TOP1cc by TOP1 poisons generates deleterious single-strand breaks (SSBs).^14^ Collision between the replication fork and TOP1cc produces DNA double-strand breaks (DSBs) and triggers cell death.^15^ The lethal TOP1cc can be trapped by endogenous DNA lesions and oxidative stress induced base modifications.^13, 16^ Therefore, the irreversible TOP1cc must be countered by endogenous repair mechanism to preserve genome integrity.^13, 17^ Tyrosyl-DNA phosphodiesterase 1 (TDP1) liberates TOP1 peptide from the 3’ end of DNA by hydrolyzing the phosphotyrosyl-bond after proteasome-dependent degradation of TOP1cc.^18^

Human TDP1, a member of the phospholipase D superfamily, has two crucial catalytic residues, H263 and H493.^19^ A homozygous TDP1 H493R mutation is associated with familial spinocerebellar ataxia with axonal neuropathy (SCAN1).^20, 21^ Lymphoblastoid cells derived from SCAN1 patients are hypersensitive to CPTs resulting from a 25-fold reduction in TDP1 activity,^22^ implying that TDP1 activity may suppress the effectiveness of CPTs. Moreover, TDP1 deficiency in *TDP1* knockdown cells confers enhanced cytotoxic response to CPT, whereas depletion of TOP1 rescues cells from CPT-induced cell death.^23, 24^ Additionally, phosphorylation of the TOP1 conserved-core domain elevates TOP1 activity, which in turn reduces cellular resistance to CPT.^25, 26^ The activity levels of both TOP1 and TDP1 may affect the anticancer efficacy of TOP1 poisons.

In this study, we measured the protein concentrations and activities of TOP1 and TDP1 in nine GBM cell lines. Our results show that the TDP1/TOP1 activity ratio has a strong correlation with irinotecan sensitivity. Furthermore, increased TDP1 activity by ectopic expression of TDP1 in GBM cells conferred increased irinotecan resistance.

## Materials and Methods

### GBM cell lines

GBM cell lines U87, A172, and H4 were obtained from American Type Culture Collection. Cell lines SF295, SF268, SF539, SNB75, SNB19, and U251 were provided by NCI Division of Cancer Treatment and Diagnosis. Cells were cultured in Dulbecco’s Modified Eagle’s medium (Corning) supplemented with 10% heat-inactivated Fetal Bovine Serum (Hyclone), 1% Penicillin/Streptomycin (Gibco), and 0.1% Plasmocin Prophylactic (InvivoGen) at 37°C in a humidified incubator with 5% CO_2_. Human astrocytes were purchased from ScienCell Research Laboratories and cultivated according to manufacturer’s protocol using Astrocyte Medium (ScienCell). Cells were used for only two subcultures to avoid senescence.

### GBM whole cell extract (WCE) preparation

To prepare WCEs, cells were plated at a density of 4 × 10^5^ cells in 60-mm dishes and collected at 80% confluence. Cells were washed twice in phosphate-buffered saline (Gibco) and lysed in radioimmunoprecipitation assay buffer (RIPA, 50 mM Tris-HCl, pH 7.4, 150 mM NaCl, 5 mM EDTA, 1 mM EGTA. 1% NP-40, 0.1% SDS and 0.5% sodium deoxycholate) supplemented with 1 mM phenylmethanesulfonyl fluoride, 1% (v/v) of Halt Protease inhibitor cocktails (Thermo Fisher), and 1% (v/v) Halt Phosphatase inhibitor cocktails (Thermo Fisher). After 5 min on ice, lyzed cells were transferred to a sterile microcentrifuge tube, followed by 30%, 30 s sonication with a Fisher Scientific 120W microtip sonicator. The cell debris was removed by centrifugation at 15,000 x g at 4°C for 15 min and supernatant protein concentration was measured using Pierce BCA Assay Kit.

### Western blots

Protein expression levels were measured using Western blots as described in the Methods Supplement. The primary and secondary antibodies are listed in Supplementary Table S1. The target protein images were obtained with C-DiGit Blot scanner (LI-COR) and analyzed by ImageStudio (LI-COR). The TOP1 or TDP1 signal intensity for each WCE was first normalized to actin signal to correct for loading, then divided by the NHA (fetal normal human astrocyte) TOP1 or TDP1 signal intensity for comparison of relative TOP1 or TDP1 protein levels. GraphPad Prism version 8.1 was used for the data analysis. The mean and standard deviations shown in bar graphs were calculated based on at least 3 technical replicates. Degree of correlations between two parameters tested was determined by using the Pearson correlation coefficient value (R) and considered as significant for P < 0.05. Each entire set of experiments were repeated three times.

### Irinotecan IC_50_ measurement for GBM cell lines

Cells were plated at a density of 8.5 × 10^3^ cells/well in a 96-well plate and cultured until 70-80% confluence. Irinotecan dissolved in DMSO was serially diluted (0.05 to 15 µM) in freshly prepared media for 72 h further incubation with the cells. The cell viabilities were assayed by TO-PRO3 (Thermo Fisher) according to the manufacturer’s protocol. The fluorescence signal from TO-PRO3 reagent was detected by Odyssey CLx Imaging System (LI-COR) from surviving cells and analyzed by ImageStudio (LI-COR). IC_50_ is determined as the irinotecan concentration resulting in 50% of cell viability compared to DMSO control. Results were obtained from triplicated experiments.

### Site-direct mutagenesis of TDP1 cDNA clone

The TDP1 cDNA clone with expression under the control of the CMV promoter in pCMV6-XL4 was obtained from OriGene. H263A substitution was performed using the Q5 Site-Directed Mutagenesis Kit (NEB) with 5’-TGGAACACACGCCACGAAAATGATG-3’ and 5’-AAC GCAATATCCAACTTTG-3’ primers to generate a mutant TDP1 clone with null catalytic activity.^19^ The complete TDP1 coding sequence in wild-type (WT) and H263A mutant clones were verified by DNA sequencing.

### Transfection with TDP1 cDNA Clones

The transient transfection of GBM cell line H4, with WT-TDP1 and H263A-TDP1 clones were performed with Lipofectamine 3000 (Invitrogen) according to its manufacturer’s protocol. Briefly, cells were seeded at a density of 1.8 × 10^5^ cells/well in 35-mm dishes and cultured 24 h until 60% confluence. DNA (500 ng) was mixed with Lipofectamine 3000 at a ratio of 1:2 (w/v) in 250 µL of Opti-MEM Medium (Gibco) for 20 min at RT and added to cells. After further incubation for 24 h, WCEs were prepared using RIPA buffer as described above.

For measurement of irinotecan IC_50_, H4 cells were plated at a density of 6.5 × 10^3^ cells/well and cultured 24 h until 60% confluence. DNA was mixed with Lipofectamine 3000 in 10 µL of Opti-MEM Medium for 20 min at RT and incubated with cells for 24 h before the addition of serial dilutions of irinotecan. After further incubation for 72 h, cell viabilities were measured with the TO-PRO3 assay as described above.

### Collection and storage of glioblastoma tumor samples from patients

Glioblastoma tumors were collected from the patients during operations at Miami Neuroscience Center of Larkin Hospital according to the protocol approved by Florida International University Institutional Review Board (FIU IRB, No. IRB-16-0355-CR02), and snap-frozen in cryotubes pre-cooled on dry ice. Samples kept in dry ice were then delivered by courier to FIU and cryopreserved in liquid nitrogen.

### Protein extraction from GBM patient samples

Ten GBM tumors from patients (de-identified data shown in Supplementary Table S2) were used to prepare WCEs. Frozen tumor tissues were transferred onto an ice-cooled 100-mm dish and sliced into small pieces. The tumor segments were then weighed and RIPA/inhibitors lysis buffer was added at a ratio of 1:10 (v/w). Samples were homogenized by Dounce Homogenizer and rotated at 4°C for 2 h followed by centrifugation at 15000 x g for 15 min. Supernatant was kept in aliquots at -80°C for further analysis.

### TOP1 relaxation activity measurement

Negatively supercoiled pBAD/Thio plasmid DNA (240 ng) purified by CsCl gradient centrifugation was used as substrate for the TOP1 relaxation assay. The reaction buffer contained EDTA to suppress the activities from the metal ion-dependent topoisomerase and nuclease activities present in the WCEs. The reaction was carried out in 20 µL of 10 mM Tris-HCl, pH 7.9, 1 mM EDTA, 150 mM NaCl, 0.1% BSA, 0.1 mM spermidine, 5% glycerol. After 30 min at 37°C, reactions were stopped with 4 µL of 6% SDS, 0.3% bromophenol blue, 30% glycerol, and analyzed by agarose gel electrophoresis.^27^ DNA in the gel was stained with EtBr and photographed over UV light. AlphaView (ProteinSimple) was used to analyze the fraction of supercoiled DNA substrate converted into relaxed DNA. Serial dilutions of recombinant TOP1 (TopoGEN) was used to generate a reference standard curve for fraction of supercoiled DNA converted to relaxed DNA by units (U) of TOP1 as defined by the supplier. Serial dilutions of WCEs were assayed under the same conditions to identify amount needed to relax 50% of the supercoiled DNA substrate and calculate the TOP1 activity present in each WCE based on the standard curve as U/µg of total WCE protein. The relative TOP1 activity levels present in the cell lines were normalized by the TOP1 activity level in NHA.

### TDP1 activity measurement by gel-based assay

The P12Y oligonucleotide linked to tyrosine at the 3’-end (5’-HO-GAAAAAAGAGTT-PO_4_-Tyr-3’, TopoGEN) was labeled at the 5’-end with ^32^P. Recombinant human TDP1 was purified as described.^28^ Serial dilutions of TDP1 was incubated with P12Y substrate (4 ng) at 37°C for 30 min in 5 µL of 20 mM Tris-HCl, pH 7.5, 100 mM KCl, 10 mM EDTA, 1 mM DTT. Reaction was stopped with 5 µL of 96% formamide, 20 mM EDTA, 0.03% xylene cyanol and 0.03% bromophenol blue, followed by heat-inactivation at 95°C for 5 min. P12Y substrate and P12 product with 3’-tyrosine removed by TDP1 were separated by electrophoresis in 20% urea-denaturing sequencing gel. Following analysis with the BioRad Pharos FX Plus Phosphorimager, the correlation between TDP1 activity, represented as fmol, and the fraction of P12 produced is plotted as a standard curve. TDP1 activity assay for WCEs was carried out under the same conditions with 2 µg of WCE proteins added to each reaction. The TDP1 activity present in each microgram of WCE was calculated based on the standard curve as fmol/µg. The TDP1 activities in the cell lines were normalized by TDP1 activity in NHA.

### TDP1 activity measurement by fluorescence-based assay

TDP1 activity was also carried out by fluorescence-based assay as described.^29^ The 5’-phosphorothioate bond linked fluorophore, FAM, and 3’-phosphodiester bond linked quencher, BHQ1, were modifications present on a 55-nucleotides long 5’FAM-DNA-BHQ1-3’ oligonucleotide substrate (5’-FAM-AAAGCAGGCTTCAACGCAACTGTGAAGATCGCTTGGGTGCGTTGAAGCCTGCTTT-BHQ1-3’, LGC BioSearch) that forms a hairpin structure. Briefly, 25 pmol of 5’FAM-DNA-BHQ1-3’ was incubated with serial dilutions of TDP1 at 37°C in 25 µL of 20 mM Tris-HCl, pH 8.0, 100 mM KCl, 10 mM EDTA, 10 mM DTT, 0.05% TritonX-100. Fluorescence signal was measured every 30 s for 500 min. The correlation between amount of TDP1, present as fmol, and the initial linear slope (0-125 min) from each reaction is plotted as a standard curve. TDP1 activity in 2 µg of WCE was assayed under the same conditions and calculated based on the standard curve as fmol/µg WCE. The relative TDP1 activity levels in the cell lines were normalized by the TDP1 activity level in NHA.

### Carboxylesterase 2 activity measurement

The carboxylesterase 2 (CES2) activity in each cell line was assayed as described^30^ using 6.7 µg of WCE proteins in the presence or absence of 300 µM of CES2 inhibitor loperamide (LOP, Sigma) in 20 µL of 100 mM Tris-HCl, pH 7.4 pre-incubated at 37°C for 10 min. Reaction was initiated by adding 200 µL of p-NPA (Sigma) substrate. Absorbance at 405 nm was recorded at 0, 5, 10, 20, 30, 40, 50, and 60 min. The amount of the reaction product p-NP formed was calculated based on the standard curve from serial dilutions of p-NP (Sigma). The carboxylesterase activity from each microgram of WCE were represented as pmol/min/µg. The difference in activity obtained in the presence and absence of LOP was considered as CES2 activity. The assay was performed in total of nine replicates in two experiments.

## Results

### Comparison of TOP1, TDP1 expression in GBM cell lines and correlation with irinotecan IC_50_

The TOP1 and TDP1 protein expression levels in GBM cell lines were compared by Western blot analysis of WCEs and normalized against NHA (Fig. 1; Supplementary Table S4). The irinotecan IC_50_ for individual cell line was measured using a cell-based assay to detect cell abundance (Supplementary Table S3). No significant correlation (R=-0.254) was found between TOP1 protein levels and irinotecan IC_50_s (Supplementary Table S9). A weak correlation between higher TDP1 protein levels and increased IC_50_ values was observed (R=0.797, P=0.01; Supplementary Table S9). These trends held across three distinct biological replicates (Supplementary Fig. S1, Table S9) suggesting that TOP1 and TDP1 levels alone may not be highly predictive indicators of irinotecan vulnerability.

**Fig 1.**
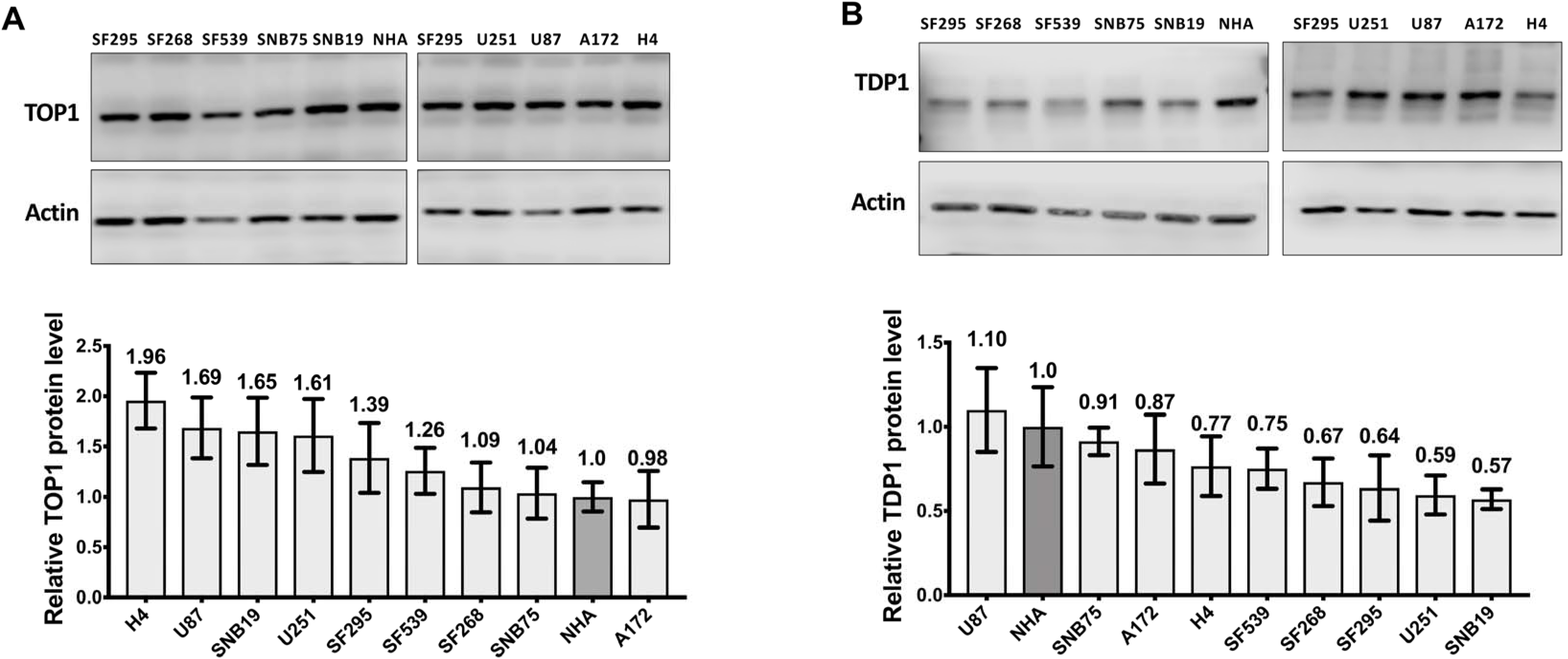
TOP1 and TDP1 protein levels in GBM cell lines. Relative levels of (A) TOP1 and (B) TDP1 protein expression in GBM cell lines were compared by Western blotting against NHA. Signal from actin was used to control loading of WCE proteins (10 µg). The average and standard deviation of measurements from one representative experiment is shown here in the bar graphs.

### TOP1 enzymatic activity in GBM cell lines does not correspond to the TOP1 protein level

Recombinant TOP1 was used to generate a linear standard curve for the increase in fraction of DNA relaxed with increasing TOP1 activity (Fig. 2A). The TOP1 activity in serial dilutions of each GBM WCE was assayed (Fig. 2B) and normalized against NHA (Supplementary Table S5). Results from additional biological replicates are shown in Supplementary Fig. S2. The relative TOP1 activity levels did not correlate with TOP1 protein concentrations (R=0.446, Supplementary Fig. S2 and Table S13) suggesting that TOP1 activity in GBM might be modulated by other means. Also, cell specific TOP1 activities did not correlate with irinotecan IC_50_ (R<0.7, Supplementary Table S9). These experiments indicate that TOP1 activity cannot predict irinotecan cytotoxicity in GBM cell lines.

**Figure 2.**
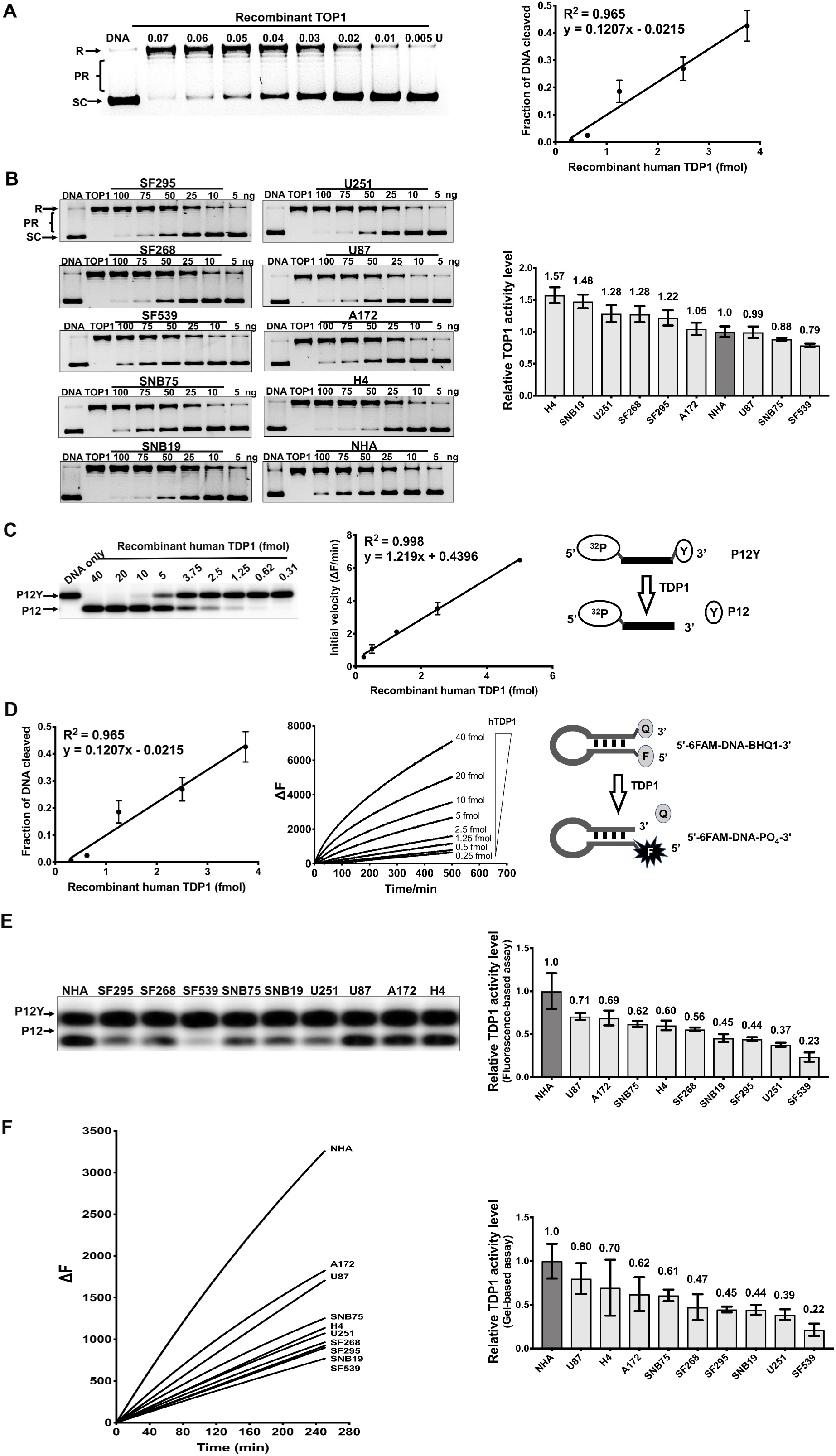
TOP1 and TDP1 activities in GBM cell lines. (A) Supercoiled plasmid DNA was relaxed by serially diluted recombinant TOP1 and DNA topoisomers were separated by gel electrophoresis. SC: supercoiled, R: relaxed, PR: partially relaxed DNA. Fraction of DNA relaxed by TOP1 was quantitated to generate a standard curve. (B) Assay of TOP1 activity in serial dilutions of GBM WCEs. The TOP1 activity levels relative to NHA are shown in the bar graph. (C) Gel assay for conversion of radiolabeled P12Y DNA substrate into P12 product by recombinant TDP1. The standard curve for fraction of substrate cleaved by increasing amounts of TDP1 is shown in the middle of the panel. (D) TDP1 3’-endonuclease activity on hairpin substrate with 5’-fluorophore and 3’-quencher leads to increase in fluorescence. The initial velocity of fluorescence increase was used to generate a standard curve. (E) Gel-based and (F) Fluorescence-based assay of relative TDP1 activity levels in GBM WCEs.

### Measurement of TDP1 activity by gel-based and fluorescence-based assays

TDP1 can act as a broad-spectrum 3’-exonuclease to hydrolyze various phosphodiester linkages at the 3’-ends of DNA.^19, 31^ A gel-based assay measured TDP1 activity with a 5’-^32^P labeled oligonucleotide with a 3’-tyrosine modification (P12Y) as substrate to yield P12 as the TDP1 reaction product in proportion to increasing amount of recombinant TDP1 (Fig. 2C). In an alternative fluorescence-based assay, a hairpin-structured oligonucleotide (Fig. 2D) with a 5’ fluorophore and a 3’-BHQ1 modification was used as a substrate.^29^ TDP1 removal of the quencher at the 3’-end leads to an increase in fluorescence signal (ΔF). Activities in the GBM WCEs measured in triplicates by the gel-based (Fig. 2E) or fluorescence-based assay (Fig. 2F) were calculated based on the standard curve of recombinant TDP1 and normalized to TDP1 activity level in NHA (Supplementary Table S6). Results from additional replicates are shown in Supplementary Fig. S3. The TDP1 activities obtained from the gel-based and fluorescence-based assays showed strong correlation (R = 0.943, Supplementary Fig. S3, panel D4), demonstrating their similar capabilities for measuring TDP1 activities *in vitro*. TDP1 activity levels assayed by the two methods correlate only slightly with TDP1 protein levels measured by Western blot (Supplementary Table S13) and irinotecan IC_50_ (Supplementary Table S9).

### TDP1/TOP1 activity ratio in GBM WCE is a strong predictor of irinotecan IC_50_

The lack of correlation between TOP1 and TDP1 protein expression (Supplementary Table S7) suggests that there is no overlapping pathway for regulating TOP1 and TDP1 levels. To further identify more promising indicators for irinotecan sensitivity, we calculated the TDP1/TOP1 protein level ratio and the TDP1/TOP1 activity level ratio (Supplementary Table S8). The TDP1/TOP1 protein ratios (Fig. 3A) were weakly correlated with irinotecan IC_50_ (R=0.696, P=0.037; Fig. 3B). However, the TDP1/TOP1 activity ratios (Fig. 3C) were strongly correlated with the irinotecan IC_50_ values (R=0.917, P=0.0005 for gel-based TDP1 assay; R=0.922, P=0.0004 for fluorescence-based TDP1 assay; Fig. 3D). The results from two other sets of replicated experiments also confirmed that the TDP1/TOP1 activity ratio is a strong predictor for irinotecan IC_50_ (Supplementary Table S9).

**Figure 3.**
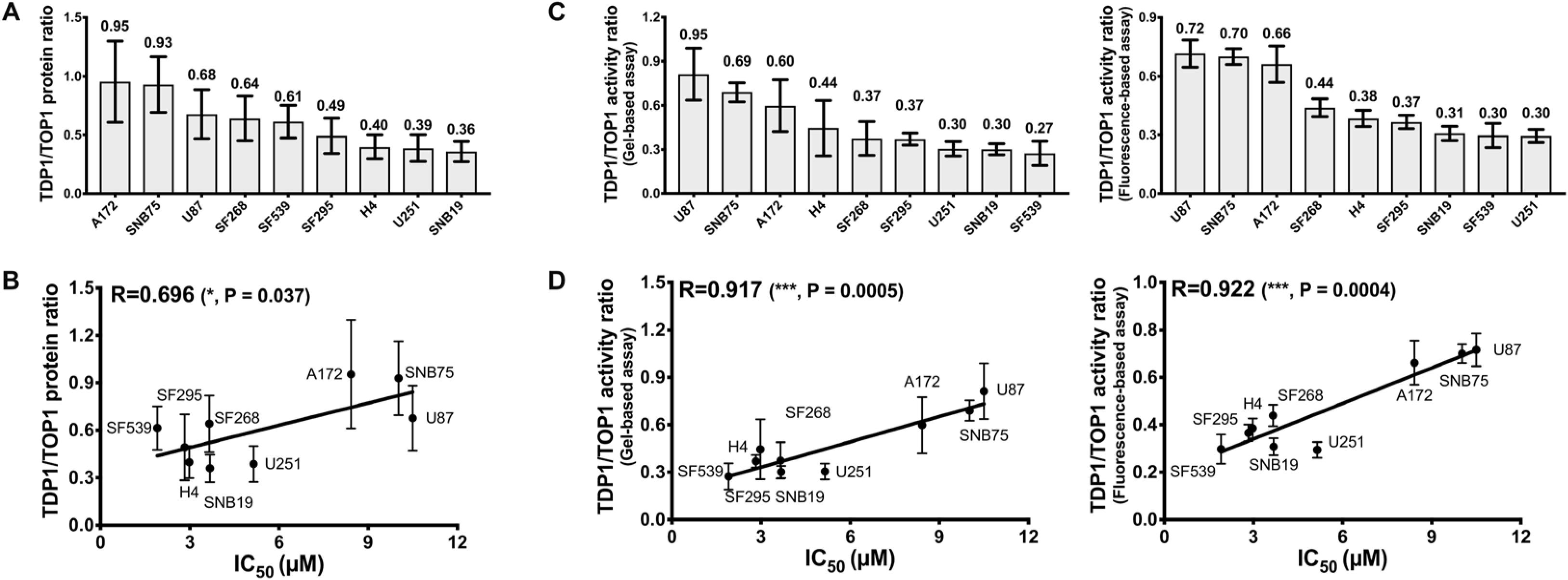
TDP1/TOP1 activity ratio is a strong predictor for irinotecan IC_50_. (A) TDP1/TOP1 protein ratio and (C) TDP1/TOP1 activity ratio in GBM WCEs. Pearson correlation between (B) TDP1/TOP1 protein ratio or (D) TDP1/TOP1 activity ratio with irinotecan IC_50_ for GBM cell lines.

### Increased resistance to irinotecan following transfection with recombinant TDP1

To follow up on the observed correlation between irinotecan sensitivity and TDP1/TOP1 activity ratio, we elevated the TDP1 activity in H4 cell line by transfection with clones expressing WT-TDP1 or null activity mutant H263A-TDP119 and determined the effect on irinotecan IC_50_. Overexpression of TDP1 and unchanged expression of TOP1 were confirmed by Western blotting (Fig. 4A). Increase in TDP1 catalytic activity in the WCE from WT-TDP1 but not H263A-TDP1 was observed (Fig. 4B) along with no change in TOP1 catalytic activity as expected. H4 cells transfected with WT-TDP1 had significantly higher Irinotecan IC_50_ than H4 cells transfected with H236A-TDP1 clone (Fig. 4C). This result further supports the correlation between TDP1/TOP1 activity ratio and irinotecan IC_50_ for GBM.

**Figure 4.**
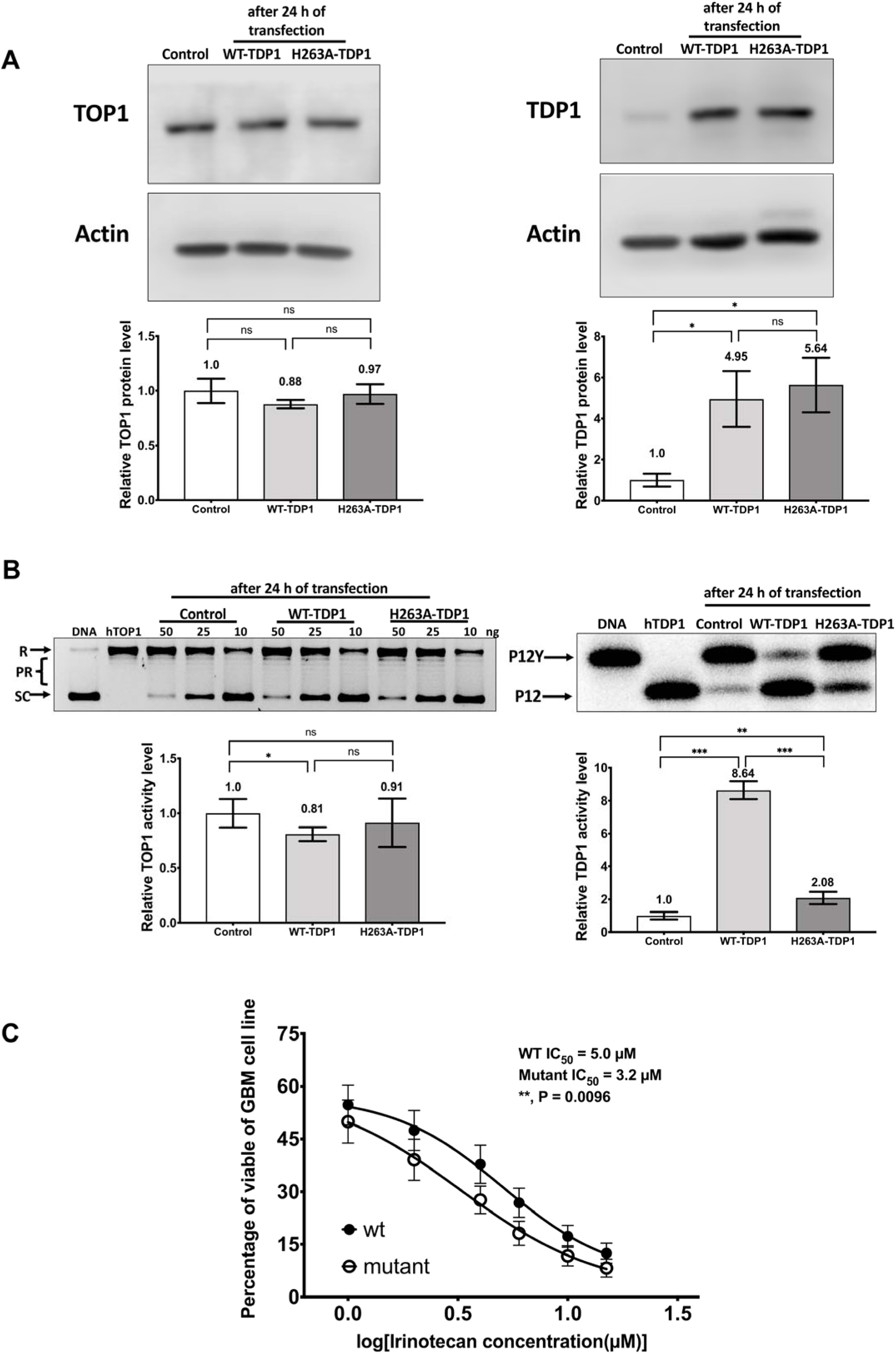
Transfection of TDP1 into H4 cell line elevates the TDP1/TOP1 activity ratio and increases the irinotecan IC_50_. (A) TDP1 and TOP1 protein levels after transfection of H4 with WT or H263-mutant TDP1 clones. (B) TDP1 and TOP1 activity levels in H4 WCE from transfection of WT-TDP1. (C) Irinotecan IC_50_ for H4 cells transfected with WT-TDP1 clone versus H263 mutant TDP1 clone (n=7).

### Carboxylesterase 2 (CES2) activity in the GBM cell lines

Irinotecan is converted to the much more potent metabolite SN-38 by carboxylesterase isoform CES2.^32^ The CES2 activity present in the GBM cell lines were assayed and found to be comparable (Supplementary Fig. S5). This demonstrates that the relatively greater irinotecan resistance for cell lines such as A172 and SNB75 was not due to lack of CES2 activity, and irinotecan sensitivity of SNB19 was not due to relatively high level of CES2 being present in SNB19. Metabolic conversion of irinotecan to SN-38 could be significant for irinotecan treatment efficacy *in vivo*.^33^ Nevertheless, the observation made here that GBM cells with lower TDP1/TOP1 activity ratio would be more sensitive to TOP1 poison is valid with the GBM CES2 activity data being taken into consideration.

### Variable levels of TDP1, TOP1 activities and TDP1/TOP1 activity ratio in GBM patient tumor cell lysates

TOP1 and TDP1 expression as well as activity in the WCEs of ten GBM patient tumors were measured (Fig. 5). The results (Supplementary Table S10) showed that the GBM patient tumor WCEs exhibited a greater range of TOP1 expression and activity than TDP1. The range of TDP1/TOP1 protein and activity ratio is shown in Fig. 6. It should be noted that tumor #62 is not included in the graph for TDP1/TOP1 activity ratio in Fig. 6B because of the extremely low TOP1 activity in the WCE of this tumor specimen. The results show a better correlation between the TOP1, TDP1 activity levels and TOP1, TDP1 protein levels for the GBM patient tumors than the GBM cell line (Supplementary Table S13). GBM patients with relatively high or low TDP1/TOP1 protein or activity ratio in the tumors could potentially be identified in the process of chemotherapy treatment selection.

**Figure 5.**
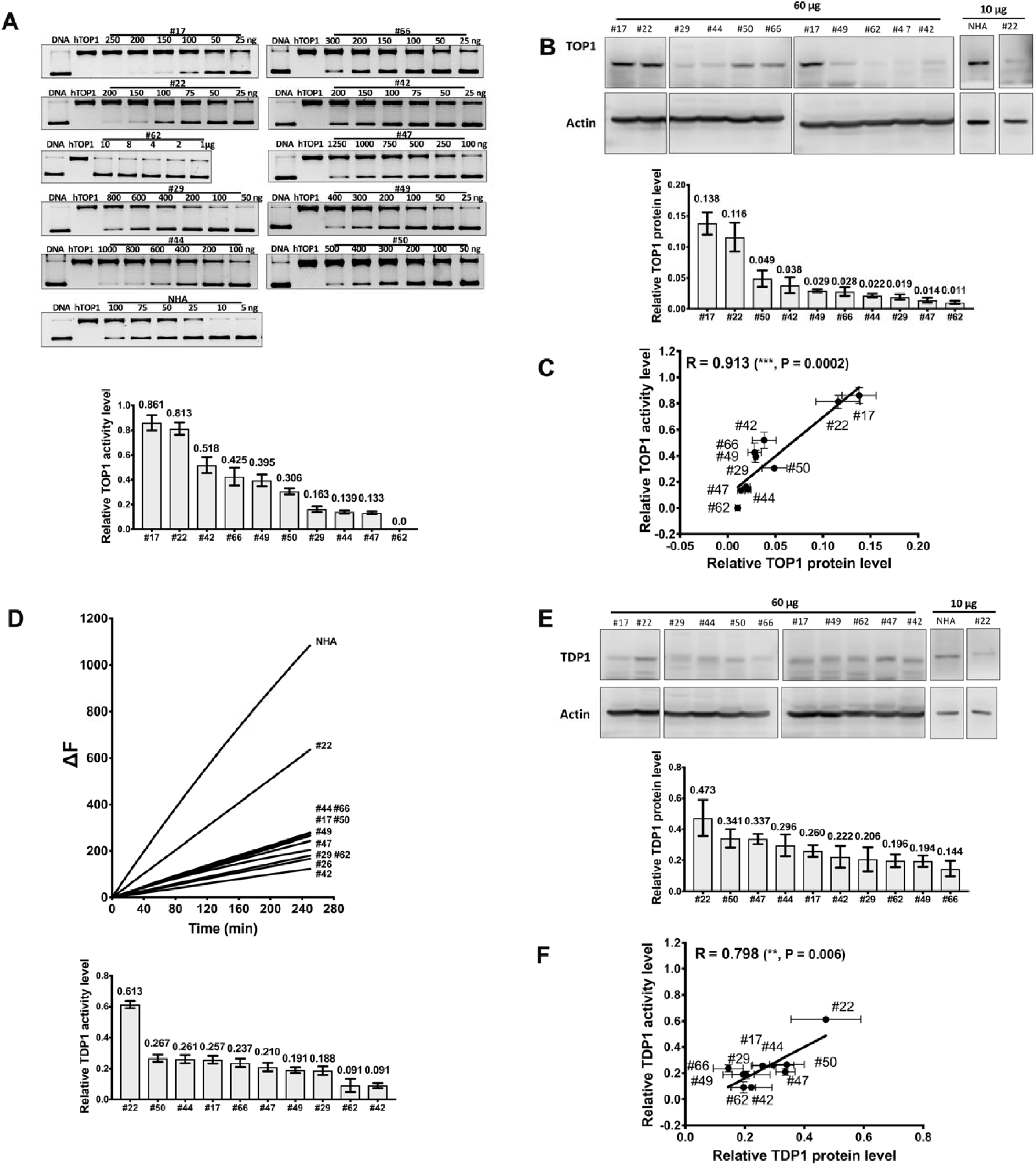
TOP1 and TDP1 activity and protein levels in GBM patient tumor WCEs. (A) Assay of TOP1 activity in GBM patient tumor WCEs. (B) Western blot analysis of TOP1 protein expression. The first two panels are from the same blot. (C) Pearson correlation of relative TOP1 protein and activity levels in GBM patient tumor WCEs. (D) Fluorescence-based assay of TDP1 activity. (E) Western blot analysis of TDP1 protein expression. (F) Pearson correlation of relative TDP1 protein and activity levels in GBM patient tumor WCEs. The bar graphs (n=3 or more) show relative TOP1 and TDP1 activity or protein levels normalized against activity and protein expression in NHA.

**Figure 6.**
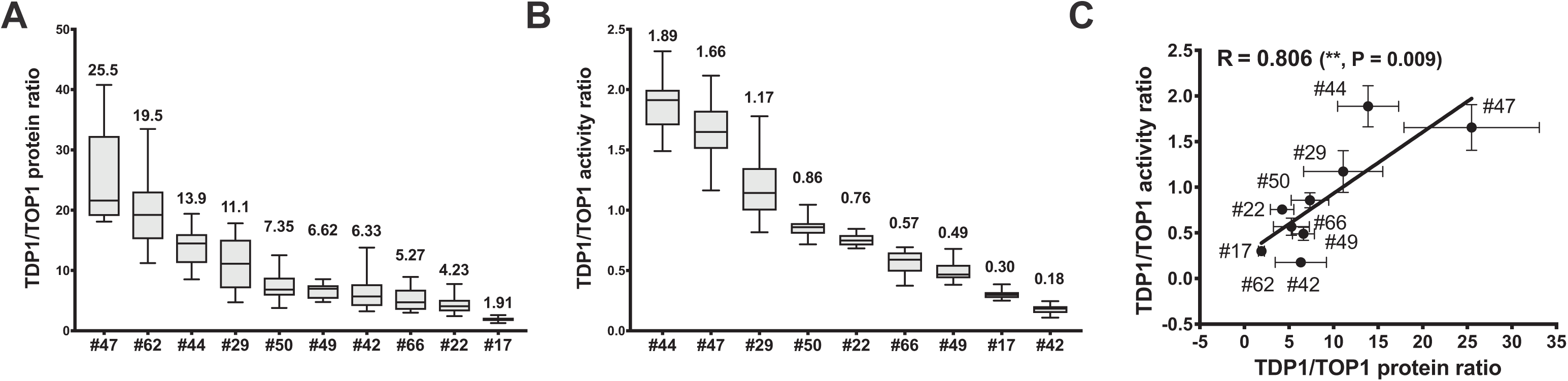
Range of TDP1/TOP1 protein ratio and activity ratios in GBM patient tumor WCEs. The box plots show the median with minimum and maximum values at the bottom and top tail-ends for (A) TDP1/TOP1protein ratio; (B) TDP/TOP1 activity ratio calculated from the relative TOP1, TDP1 protein and activity levels of each GBM patient tumor WCE. (C) Pearson correlation between the TDP1/TOP1 protein ratio and TDP1/TOP1 activity ratio.

## Discussion

GBM is a devastating disease with poor prognosis and lack of predictive biomarkers for chemotherapy.^1^ The prodrug irinotecan is converted to a topoisomerase I poison specifically to stabilize the TOP1cc on chromosomal DNA. Irinotecan has been found in clinical trials to exhibit modest and not consistently reproducible efficacy in the treatment of GBM. Therefore, our study is aimed at evaluating the roles of TOP1 as target and the TOP1cc repair enzyme TDP1 on the sensitivity of GBM to irinotecan in order to identify rational predictive biomarkers that may contribute to improving the treatment outcomes.

Previous studies have shown that cytotoxicity of TOP1 poisons is associated with TOP1 and TDP1 protein expression as well as their catalytic activities. Infection of GBM cell lines with adenovirus was found to increase TOP1 expression and activity, and enhance the antiglioma effect of irinotecan.^34^ A preliminary study on 9 clinical colorectal tumors indicated that the 3 samples with the highest TOP1 expression and activity are from patients who responded to irinotecan.^35^ A later study on the roles of TDP1 and TOP1 in modulating colorectal cancer response to irinotecan demonstrated that TDP1 overexpression or TOP1 depletion is protective while conversely TDP1 depletion leads to TOP1-dependent hypersensitivity to irinotecan, but there was no correlation between inherent TDP1 or TOP1 protein levels alone and irinotecan sensitivity.^23^ A study on SCLC cell lines found that TDP1/TOP1 protein ratio is a better predictor than the individual TDP1 or TOP1 protein level.^36^

In our study, we found that the TDP1/TOP1 activity ratio is superior to TDP1/TOP1 protein ratio as a predictor for the response of GBM cell lines to irinotecan treatment as indicated by irinotecan IC_50_s. This may be due to post-translational modification (PTM) of TOP1 and TDP1 proteins in GBM. TOP1 phosphorylation increases TOP1 activities and cellular sensitivity to CPT.^26, 37^ O-GlcNAcylation can also elevate the activity of TOP1.^38^ The PTMs could confer TOP1 activity alterations and might account for the weaker correlation between TOP1 protein and its activity levels in GBM cell lines. In contrast, the lesser influence of PTM on TDP1 activity might account for the stronger correlation between TDP1 protein and its activity level. The z-scores for irinotecan, camptothecin and topotecan are available in the CellMiner database for the 6 GBM cell lines included in the NCI-60 panel (Supplementary Table S11).^39, 40^ The sensitivity z-scores for these TOP1 inhibitors all show better correlation with our TDP1/TOP1 activity ratio than the TDP1/TOP1 protein ratio or TDP1 protein and activity level alone (Supplementary Table S12). As expected, there is no correlation between TDP1/TOP1 activity ratio and sensitivity z-scores for TOP2 inhibitors in CellMiner (Supplementary Table S12).

The increase of irinotecan IC_50_ resulting from transfection of GBM H4 cell with recombinant TDP1 clone is consistent with TDP1 activity being an important determinant of GBM sensitivity to irinotecan. Combination therapy that targets both TOP1 and TDP1 has great potential to improve the treatment outcomes of GBM. Therefore, the screening of TDP1 inhibitors is a promising venue for developing new cancer therapies.^18, 41, 42^

There are additional cellular factors that contribute to irinotecan treatment outcome. Degradation of TOP1cc is necessary for sufficient exposure of the phosphodiester linkage to TDP1 hydrolysis.^13^ Previous studies have shown that DNA damage-induced PTMs are associated with TDP1 N-terminal domain (NTD, residue 1-148). Phosphorylation, SUMOylation, and PARylation at NTD stabilize TDP1 and promote TDP1 recruitment on DNA damage sites without interfering with its activity.^43–46^

Drug sensitivities are also determined in part by alternative repair pathways and epigenetic modifications. PNKP and XRCC1 are required for SSB repair following the TDP1 repair process.^13^ RECQ1, Mus81-EME1, and XPF-ERCC1 could also provide resistance to CPTs on either SSB or DSB repair pathways.^47–49^ Perturbed histone acetylation stimulates faster access of repair enzymes to the damaged sites and subsequently a higher resistance.^50^

There is good correlation between TOP1 activity and TOP1 expression level in the ten GBM patient tumors examined here (R=0.913, Supplementary Table S13), so the use of the TDP1/TOP1 protein ratio as a predictive biomarker for irinotecan sensitivity should be further examined along with TDP1/TOP1 activity ratio for GBM patient tumors.

In summary, we demonstrate that the variation in TDP1/TOP1 activity ratio in GBM could be a potential predictive biomarker for irinotecan treatment. Further study on TDP1 and TOP1 protein expression and activity in patient derived GBM models along with clinical outcome is required.

## Acknowledgment

We acknowledge the assistance of Kristen Rothenberg at Miami Neuroscience Center at Larkin.

---

**Fig S1.**
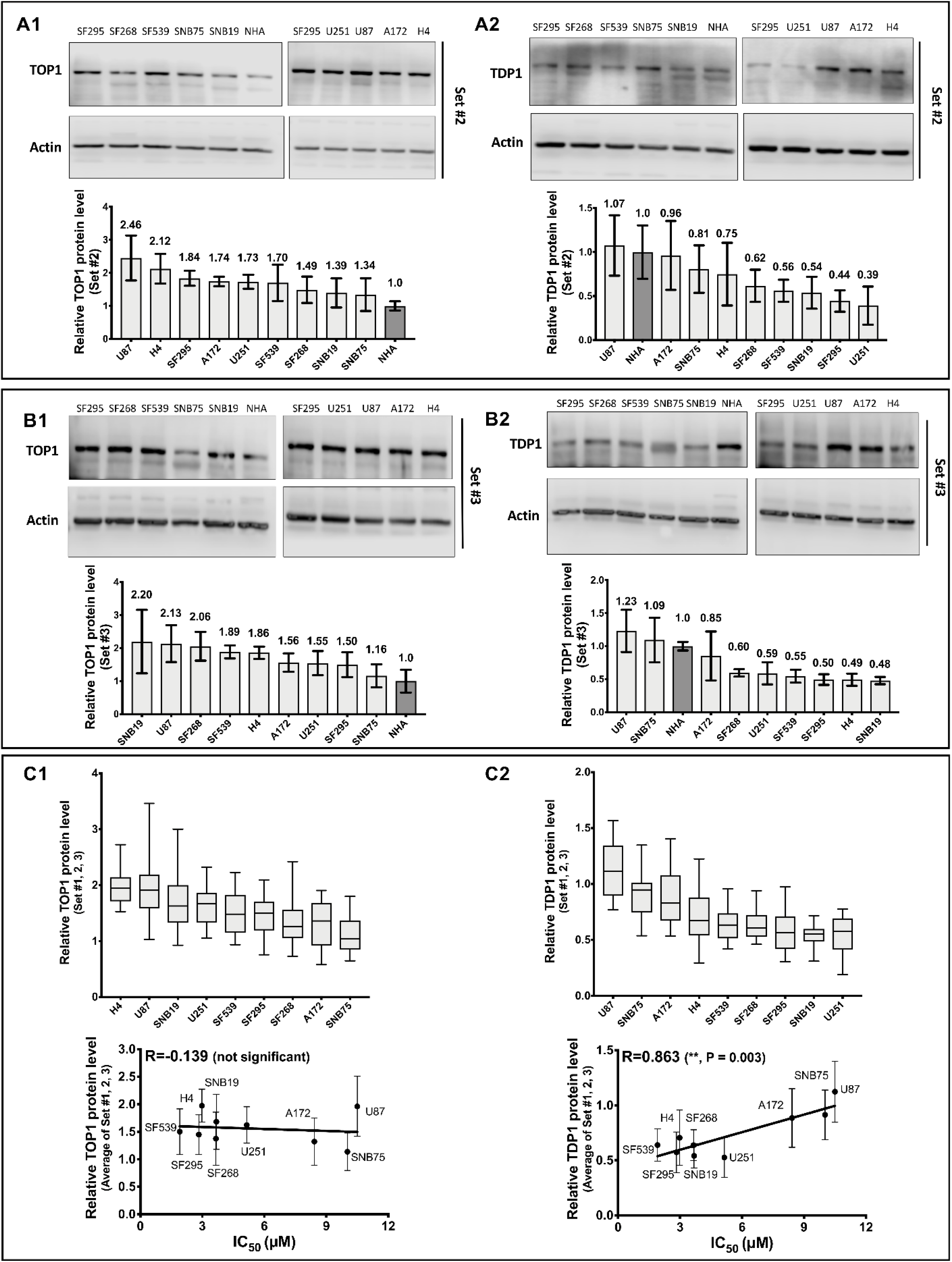
TOP1 and TDP1 protein levels measurements in different sets of GBM cell lines to assess potential correlation with irinotecan IC_50_. Relative levels of TOP1 protein expression in (A1) set #2 and (B1) set #3 GBM cell lines were compared against NHA by Western blotting. Relative levels of TDP1 protein expression in (A2) set #2 and (B2) set #3 were also compared by Western blotting against NHA. Signal from actin was used to control the loading of WCE (10 µg total protein). The bar graphs show the average and standard deviation for at least 3 measurements in each set of experiment. The box plots in (C1, C2) show the median of data from all three sets of experiments with minimum and maximum values at the bottom and top tail-ends. Pearson correlations between irinotecan IC_50_ values and relative TOP1 protein level (C1) or relative TDP1 protein levels (C2) are shown for the average of set #1, #2, #3 GBM cell lines.

**Fig S2.**
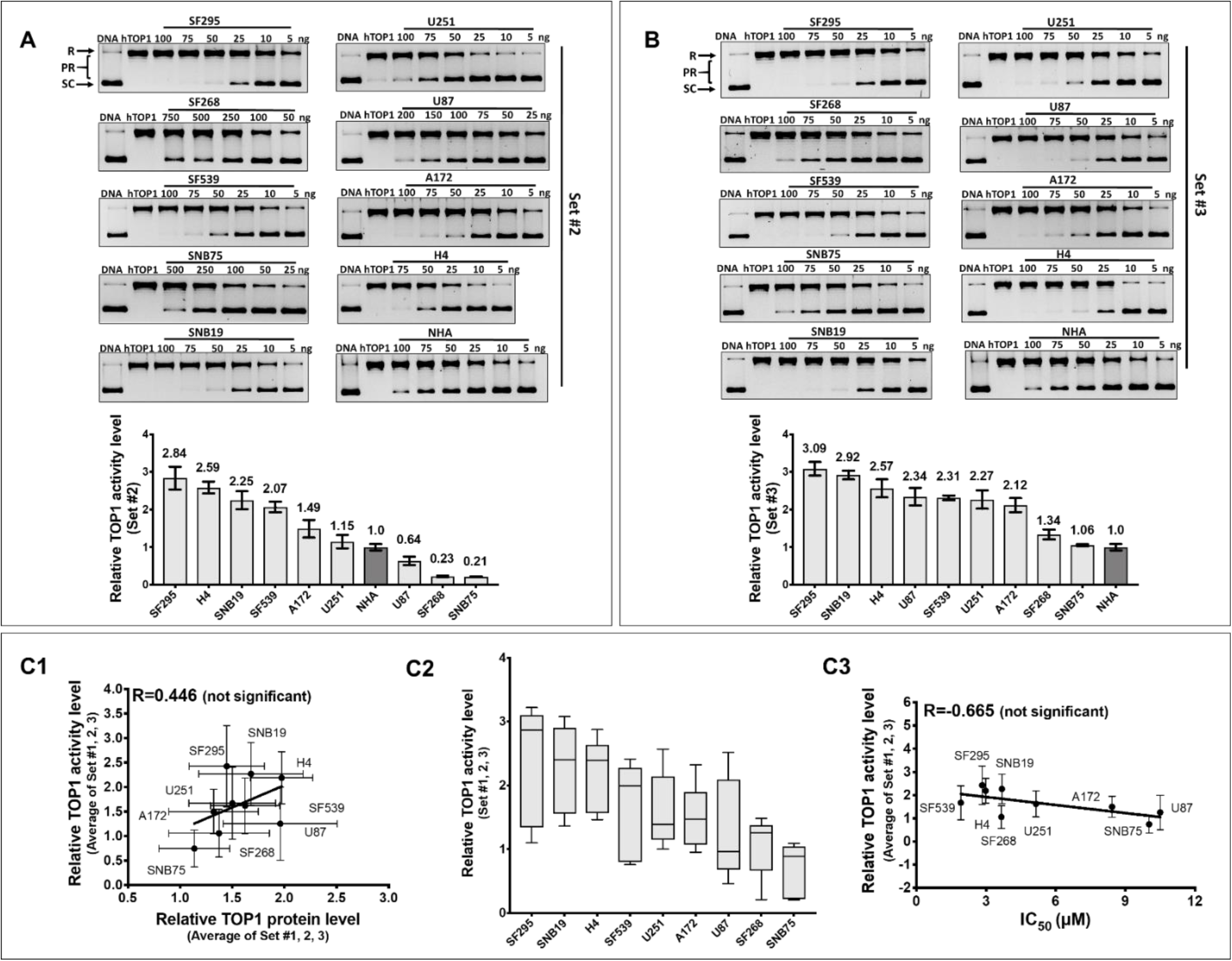
Topoisomerase I (TOP1) activity in replicated sets of GBM cell line experiments does not correlate with TOP1 protein levels or irinotecan IC_50_. Relative levels of TOP1 activity in (A) set #2 and (B) set #3 GBM cell lines were compared against NHA by plasmid DNA relaxation assay. R: relaxed DNA; PR: partially relaxed DNA; SC: supercoiled DNA. The bar graphs show the average and standard deviation for at least 3 measurements in each set of experiment. Pearson correlation between relative TOP1 protein levels and relative TOP1 activity levels are shown for the average of set #1, #2, #3 GBM cell lines (C1). The box plot in (C2) shows the median of relative TOP1 activity from all three sets of experiments with minimum and maximum values at the bottom and top tail-ends. (C3) Pearson correlation between irinotecan IC_50_ values and relative TOP1 activity levels are shown for the average of replicates set #1, #2, #3 GBM cell lines.

**Fig S3.**
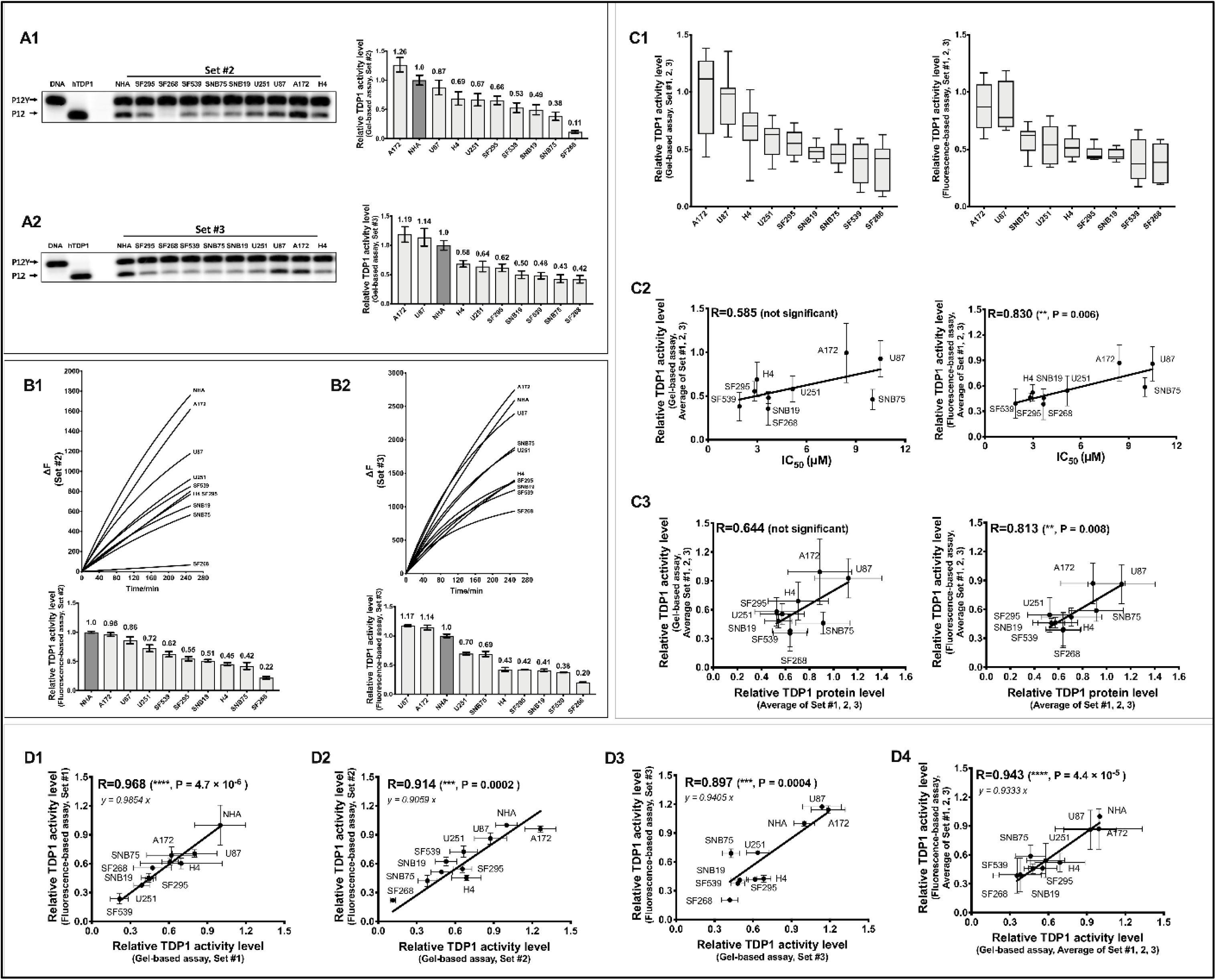
TDP1 activity levels show slight correlation with irinotecan IC_50_. Relative levels of TDP1 activity in (A1, B1) replicates set #2 and (A2, B2) set #3 were compared by gel-based assay and fluorescence-based assay against NHA. P12Y: radiolabeled DNA substrate; P12: product formed by TDP1. The bar graphs show the average and standard deviation for at least 3 measurements in each set of experiment. The box plots in (C1) show the median of relative TDP1 activity from all three sets of experiments with minimum and maximum values at the bottom and top tail-ends. Pearson correlations between the average of relative TDP1 activity from set #1, #2, #3 and irinotecan IC_50_ values are shown in (C2). Pearson correlation between relative TDP1 protein levels and TDP1 activity levels are shown in (C3) for the average from set #1, #2, #3 experiments. Pearson correlation between TDP1 activity levels measured by gel-based assay and fluorescence-based assay are shown for (D1) set #1, (D2) set #2, (D3) set #3, and (D4) average of all three sets of replicated experiments on the GBM cell lines.

**Fig S4.**
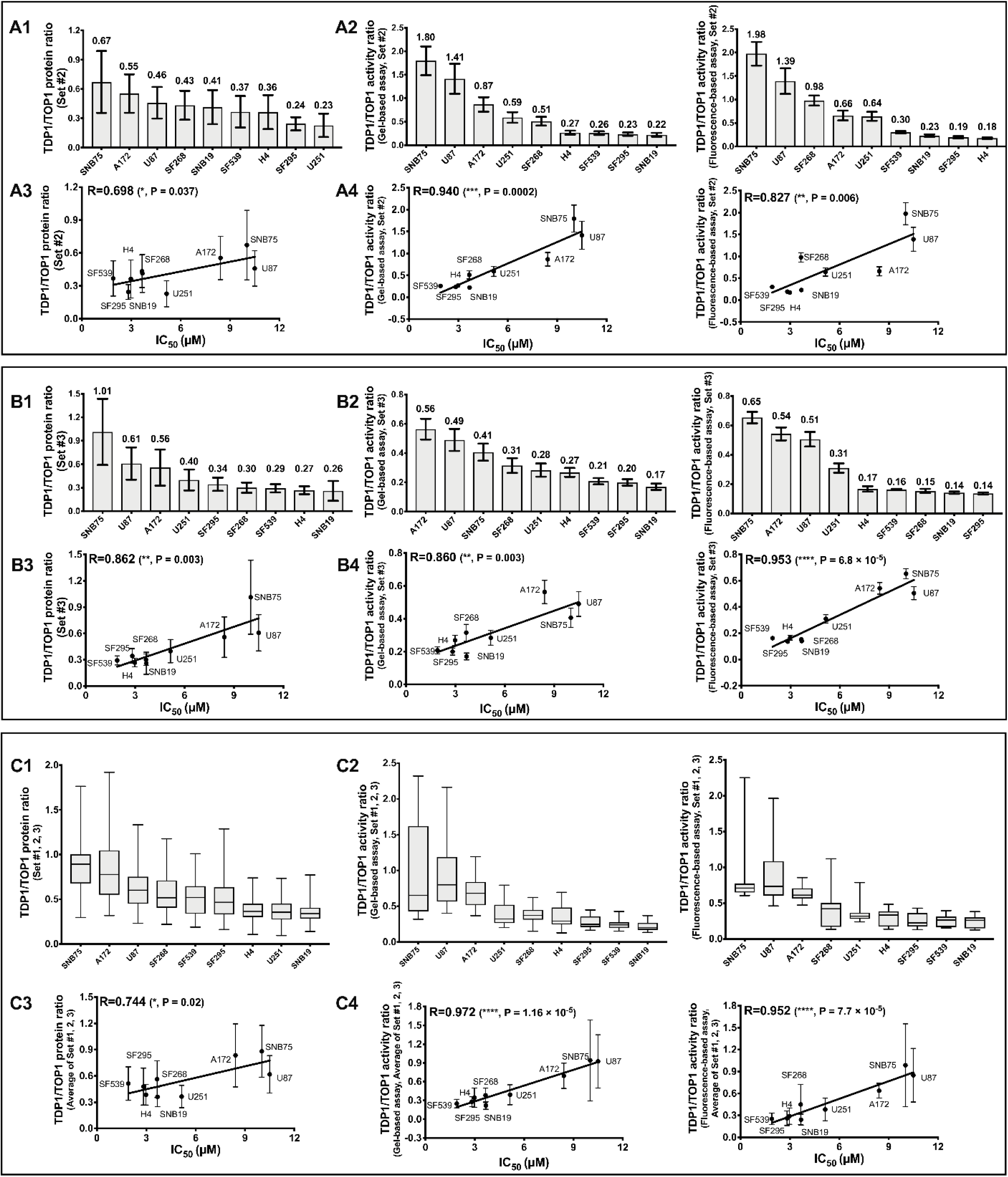
TDP1/TOP1 activity ratio is a stronger predictor than TDP1/TOP1 protein ratio for irinotecan IC_50_. Relative TDP1/TOP1 protein ratios are shown for (A1) set #2 and (B1) set #3 experiments. Relative TDP1/TOP1 activity ratios from the gel-based and fluorescence-based TDP1 activity assay are shown for (A2) set #2 and (B2) set #3. The box plots show the median of TDP1/TOP1 protein (C1) or activity ratios (C2) from all three sets of experiments with minimum and maximum values at the bottom and top tail-ends. Pearson correlations between the TDP1/TOP1 protein ratios and irinotecan IC_50_s are shown for (A3) set #2, (B3) set #3 and (C3) the average from set #1, 2, 3. Pearson correlations between the TDP1/TOP1 activity ratios and irinotecan IC_50_s are shown for (A4) set #2, (B4) set #3 and (C4) the average from replicated experiments set #1, 2, 3.

**Figure S5.**
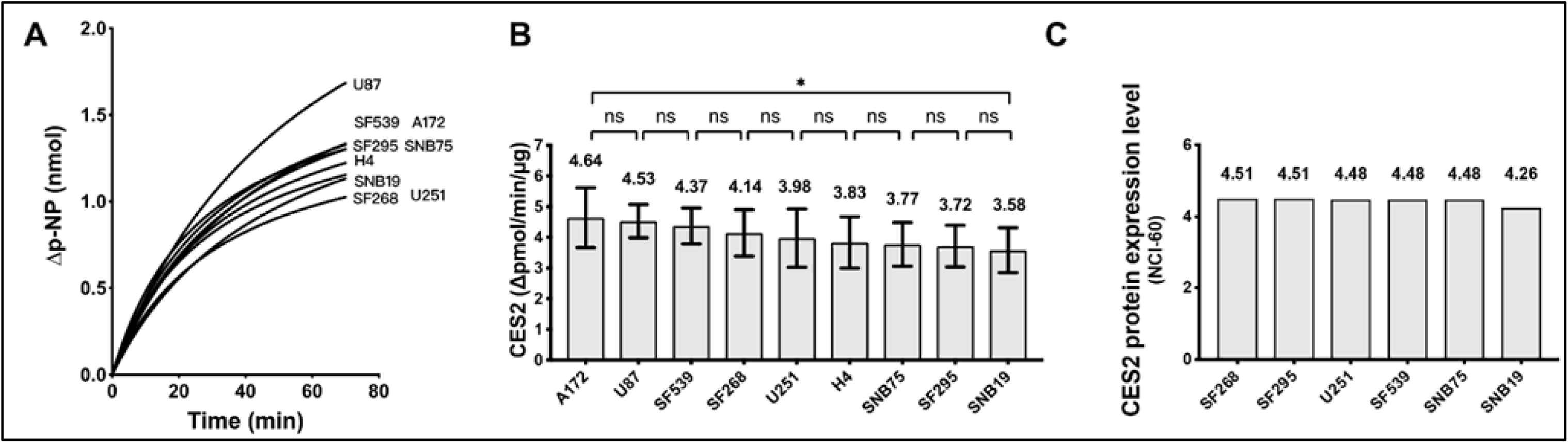
Comparison of carboxylesterase CES2 activity in WCE of GBM cell lines. (A) The time course of p-NP generation attributable to CES2 was calculated by subtracting the p-NP signal obtained in the presence of the CES2 inhibitor LOP. (B) Variation of CES2 activity in GBM cell lines measured over the initial 20 min of reaction. Results represent the average from 9 replicate measurements in two experiments. (C) CES2 protein expression levels (SWATH-MS) from the CellMinerCDB database for the GBM cell lines in the NCI-60 panel.

## Methods Supplement

**Western blot** - Proteins from whole cell extract (WCE) were first separated by 7.5% SDS-PAGE and then transferred to nitrocellulose membrane with transfer buffer (48 mM Tris, 39 mM glycine, 0.4% SDS, 20% v/v methanol) at 100 V for 1 h. The membrane was blocked with 5% bovine serum albumin (BSA) in Tris-buffered saline (TBS) at room temperature for 1 h followed by incubation of 1:1000 (v/v) primary antibody diluted in 1% Tween 20 in TBS (TBST) at 4°C for 18 h. The membrane was washed three times for 5 min (3 × 5 min) and horse-radish peroxidase (HRP) conjugated secondary antibody 1:5000 (v/v) diluted in TBST was incubated with the membrane at room temperature (RT) for additional 1 h. The membrane was then washed 3 × 5 min before the target protein expression level was developed with SuperSignal West Pico Plus Chemiluminescent Substrate (Thermo Fisher) for 5 min.

